# Autophagy in maternal tissues contributes to *Arabidopsis thaliana* seed development

**DOI:** 10.1101/2023.02.27.530228

**Authors:** Ori Erlichman, Shahar Weiss, Maria Abu-Arkia, Moria Ankary Khaner, Yoram Soroka, Weronika Jasinska, Leah Rosental, Yariv Brotman, Tamar Avin-Wittenberg

**Affiliations:** Department of Plant and Environmental Sciences, Alexander Silberman Institute of Life Sciences, The Hebrew University of Jerusalem, Givat Ram, 9190401, Jerusalem, Israel; Department of Life Sciences, Ben-Gurion University of the Negev, 8410501, Beer-Sheva, Israel

**Keywords:** *Arabidopsis thaliana*, Autophagy, Maternal tissue, Seed, Source-sink relationship, Storage compounds.

## Abstract

Seeds are an essential food source, providing nutrients for germination and early seedling growth. Degradation events in the seed and the mother plant accompany seed development. One degradation mechanism is autophagy, facilitating cellular component breakdown in the lytic organelle. Autophagy influences various aspects of plant physiology, specifically nutrient availability and remobilization, suggesting its involvement in source-sink interactions. During seed development, autophagy was shown to affect nutrient remobilization from mother plants and function in the embryo. Yet, these studies examined autophagy-knockout (*atg* mutant) plants, making it impossible to distinguish between the contribution of autophagy in the source (i.e., the mother plant) and the sink tissue (i.e., the embryo).

To address this, we employed a novel approach to differentiate between autophagy in source and sink tissues. We investigated how autophagy in the maternal tissue affects seed development by performing reciprocal crosses between WT and *atg* mutant *Arabidopsis thaliana* plants. Although F1 seedlings possessed a functional autophagy mechanism, etiolated F1 plants from maternal *atg* mutants displayed reduced growth. This was attributed to altered protein but not lipid accumulation in the seeds, suggesting autophagy differentially regulates carbon and nitrogen remobilization. Surprisingly, F1 seeds of maternal *atg* mutants exhibited faster germination, resulting from different seed coat development.

Our study emphasizes the significance of examining autophagy in a tissue-specific manner, revealing valuable insights into the interplay between different tissues during seed development. It sheds light on the tissue-specific functions of autophagy, offering potential for new research into the underlying mechanisms governing seed development and crop yield.

## Introduction

Seeds are a major source of food and feed, providing approximately 70% of the world’s human caloric intake (Sreenivasulu and Wobus, 2013). The nutritional value of seeds is attributed to storage compounds accumulating in the seed during development (Cernac and Benning, 2004). These include starch, storage lipids (mostly triacylglycerols - TAGs), and storage proteins, the relative amount of which varies between plant species. Storage compounds are degraded during germination and early seedling development to facilitate germination and seedling growth until photosynthesis establishment (Baud et al., 2008). In annual plants, seed development is accompanied by mother plant senescence, resulting in nutrient remobilization to the developing seeds, used to produce storage compounds (Distelfeld et al., 2014). Another vital seed trait is longevity, defined as seed viability after seed dry storage (Nguyen et al., 2012). Seed longevity can directly affect germination and, as a result, crop productivity and food security (Zhou et al., 2019). With a growing world population (Ort et al., 2015), specifically in light of current climate change (Lobell et al., 2011), an increased understanding of nutrient remobilization to seeds and seed longevity is vital to ensure food security.

Nutrient trafficking and partitioning to various tissues in the plant, i.e. source-sink relationship, are vital for seed development (Sonnewald and Fernie, 2018; Tegeder and Masclaux-Daubresse, 2018). Nutrients from source tissues can be used for organ growth and later exported to sink tissues following catabolism and nutrient recycling (Osorio et al., 2014; Li et al., 2015). Nutrient demand of the sink tissue can also affect export from the source (Lemoine et al., 2013). Plant source-sink relationship has been studied for many years. However, there is still a debate as to which process controls plant growth and yield the most (Sonnewald and Fernie, 2018). During seed development, both carbon, in the form of sucrose, and nitrogen, in the form of the amino acids glutamine and asparagine, are transported from the mother plant (source tissue) to the developing seeds (sink tissue) (Dourmap et al., 2023).

Macroautophagy (hereafter termed “Autophagy”) is a conserved eukaryotic mechanism for the degradation of cytoplasmic constituents in the lytic organelle (vacuoles in yeast and plants and lysosomes in animals) (Bassham et al., 2006; Marshall and Vierstra, 2018). The targets of autophagy are diverse and include long-lived proteins, protein complexes, and whole organelles (Reumann et al., 2010). The autophagy machinery relies on highly-conserved AuTophaGy-related (*ATG*) genes (Bassham, 2009; Avin-Wittenberg et al., 2012; Liu and Bassham, 2012; Ryter et al., 2013), which have been characterized in many plant species (Chung et al., 2009; Kurusu et al., 2014; Zhou et al., 2014). In recent years, autophagy was shown to be involved in many aspects of plant life, from plant development to biotic and abiotic stress response (Avin-Wittenberg et al., 2018).

Autophagy-deficient plants (*atg* mutants) are hypersensitive to carbon and nitrogen starvation and display early senescence and reduced yield (Doelling et al., 2002; Li et al., 2015; Barros et al., 2017), as well as slightly slower germination rates and impaired seedling establishment (Yoshimoto et al., 2014; Avin-Wittenberg et al., 2015). Autophagy was shown to promote nutrient remobilization in plants, functioning in the supply of lipids and primary metabolites (Masclaux-Daubresse et al., 2014; Avin-Wittenberg et al., 2015; Wada et al., 2015; Barros et al., 2017; Hirota et al., 2018; McLoughlin et al., 2018; McLoughlin et al., 2020; Barros et al., 2021). Moreover, overexpression of *ATG* genes in *Arabidopsis thaliana* (Arabidopsis) was reported to increase seed yield and fatty acid content (Minina et al., 2018).

Autophagy was shown to play a dual role in seed development, affecting both the mother plant and the developing embryo. Studies in Arabidopsis and maize (*Zea mays)* demonstrated that impaired autophagy causes altered nitrogen mobilization from the mother plant to the seed, resulting in changes in seed C/N ratio (Guiboileau et al., 2012; Li et al., 2015). Alternatively, microscopic evidence from wheat (*Triticum aestivum*) suggested the possible involvement of autophagy in the transport of storage proteins to protein storage vacuoles within the developing seeds (Levanony et al., 1992). Furthermore, an additional publication demonstrated the presence of an active autophagy mechanism within developing seeds as well as altered storage protein content and processing in Arabidopsis *atg* mutants (Di Berardino et al., 2018).

Though autophagy has been established as a major player in plant nutrient remobilization, the work thus far analyzed knockout plants, in which autophagy is inactive in both source and sink tissues. Indeed, although the studies above suggest autophagy is essential for seed development, it was impossible to distinguish between the impact of autophagy in the mother plant and in the seed itself. Moreover, the seed is comprised of several tissues with differing parental contributions. The seed coat is of maternal origin, while the embryo and endosperm are zygotic tissues, and the genomic contribution of the triploid single-layer endosperm is unevenly divided between the female parent (2n) and the male parent (1n) (Bentsink and Koornneef, 2008). In this work, we devised an experimental system that allowed us to separate the effect of autophagy in source and sink tissues and elucidate how autophagy in the maternal tissue affects seed development. To that end, we performed reciprocal crosses between Arabidopsis wild type (WT) and *atg* mutants to receive *ATG*-heterozygous F1 seeds with different maternal tissues. We discovered F1 seeds differentially accumulated storage compounds according to the genotype of the mother plant. Surprisingly, we observed that F1 seeds from maternal *atg* mutant origin had a faster germination rate, in striking contrast to homozygous *atg* mutants, which germinate slower than WT seeds. This possibly stems from differences in seed coat structure, which also influences seed longevity. Our work suggests that *atg* mutant seeds display a compound phenotype, merging the impact of autophagy in the mother plant and the embryo.

## Results

### F1 progeny of WT and *atg* mutants possess a functional autophagy mechanism but reduced growth when germinated in the dark

To differentiate between autophagy in the mother plant and in the embryo, we performed reciprocal crosses between WT and *atg* mutant (*atg5-1* or *atg7-2*) Arabidopsis plants. This novel approach allowed us to produce F1 seeds in which the embryos are heterozygous for either *atg5* or *atg7* mutation while maintaining maternal tissues that are homozygous to either WT, *atg5-1,* or *atg7-2* (Fig. 1) mutants. The heterozygous genotype of F1 plants was validated by PCR (Fig. S1a,b).

**Fig 1.**
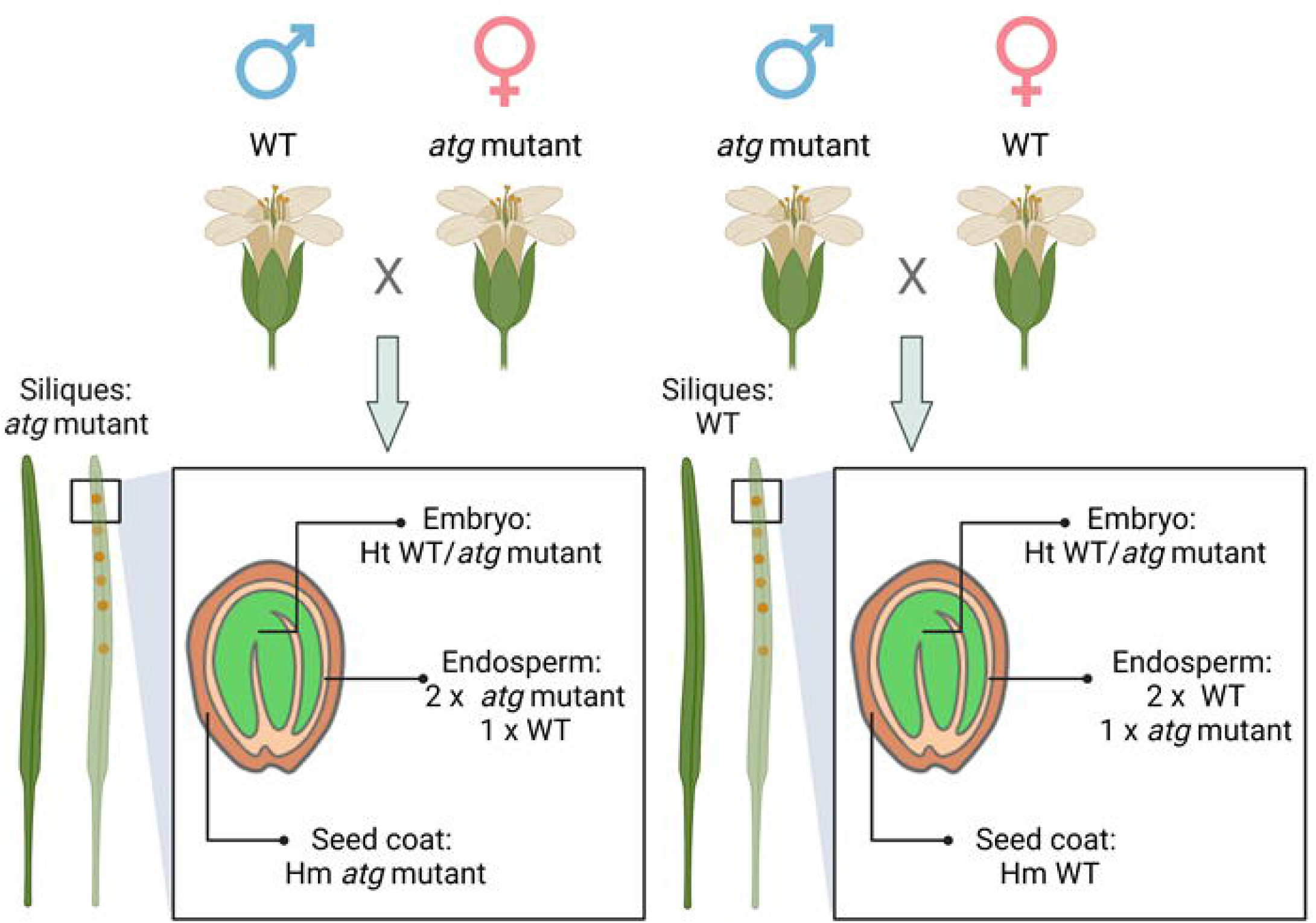
Schematic representation of the experimental system. Reciprocal crosses between WT and *atg5-1* or *atg7-2* mutants were performed. The illustration denoted the genetic makeup of the resulting F1 seeds as well as the mother plants.

We did not observe any developmental difference between the F1 seeds (Fig. 2a). In addition, we did not detect any aborted seeds, as previously reported in *atg5-1* plants (Di Berardino et al., 2018). Homozygous *atg* mutant seeds are smaller than WT seeds (Barros et al., 2017; Minina et al., 2018). We thus compared the weight of reciprocally crossed F1 seeds to examine whether we see a similar effect due to different maternal tissues. We did not see a significant difference between the lines, suggesting this phenotype might be governed by autophagy in the developing embryo (Fig. 2b).

**Fig. 2:**
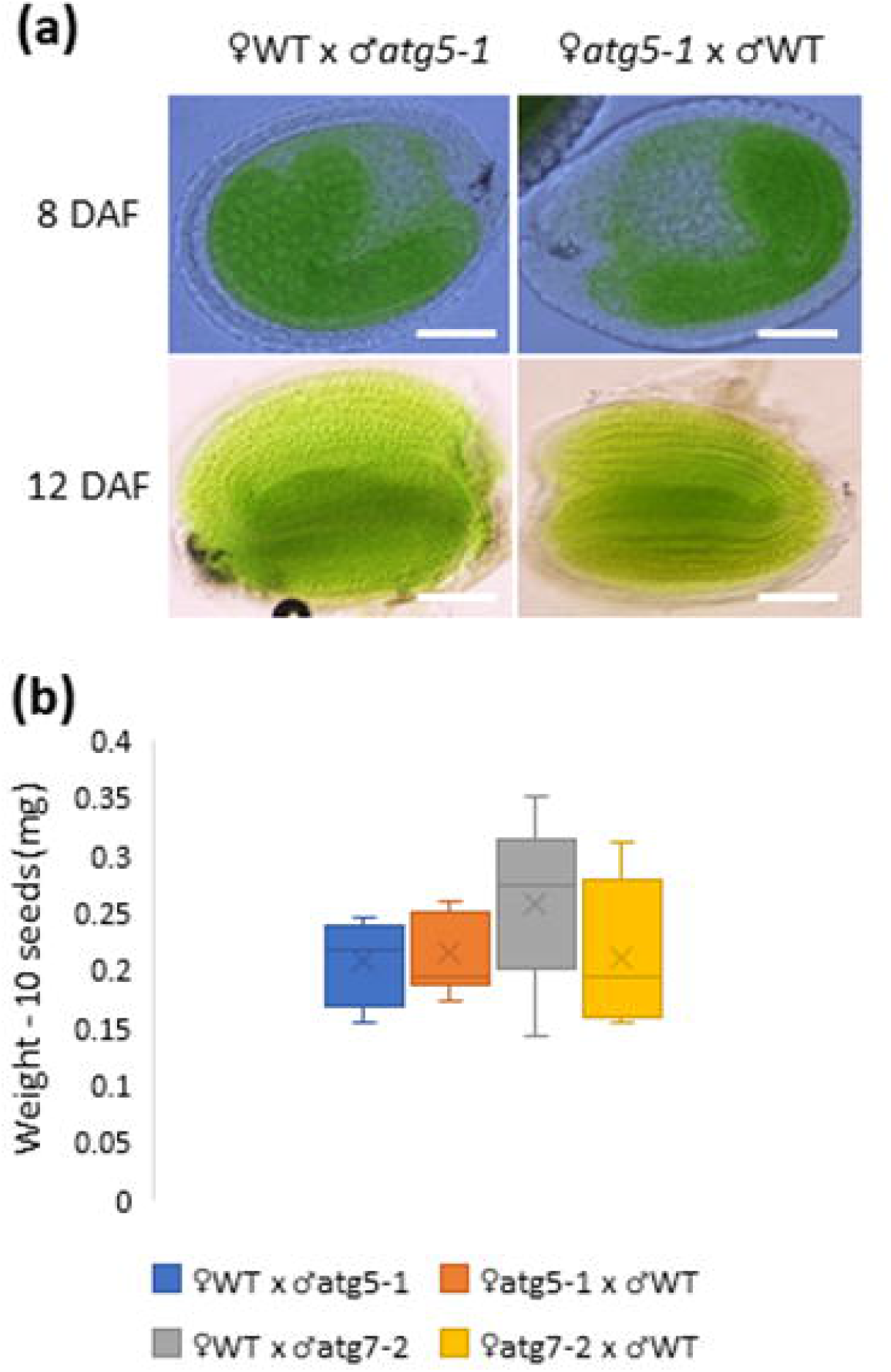
F1 progeny of WT and *atg* mutants display normal development. (a) F1 seeds were collected at 8DAF and 12DAF, uncolored, and photographed. No aborted embryos were observed. Representative image of 40±5 seeds examined. Scale bar=100µm. (b) Weight of 10 F1 seeds. Data are presented as a box&whiskers plot. No significant differences were observed (n=4-8).

We next wished to determine whether the F1 plants display a functional autophagy mechanism. One hallmark phenotype of *atg* mutants is hypersensitivity to carbon starvation (Marshall and Vierstra, 2018). We thus exposed two-week-old F1 seedlings and their respective paternal lines to carbon starvation and recovery. As can be seen in Fig. 3a-c and Fig. S1c-d, the *atg* mutant parental lines were more susceptible to carbon starvation than WT plants and did not recover, as expected. F1 seedlings displayed a WT-like phenotype, irrespective of the identity of the maternal or paternal lines. *atg* mutant etiolated seedlings were also shown to exhibit shorter hypocotyls when grown without exogenous sugars (Avin-Wittenberg et al., 2015). In light of the results of the carbon starvation experiments, we were expecting not to see a difference between the F1 progeny of our reciprocal crosses. Interestingly, when we performed this assay on the F1 seeds, seedlings stemming from *atg* mutant mother plants had significantly shorter hypocotyls than seedlings from WT mother plants (Fig. 3d,e). As etiolated seedlings are reliant on nutrients from the seeds alone, we hypothesized this phenotype resulted from altered storage compound amounts deposited in the seeds from different mother plants.

**Fig. 3:**
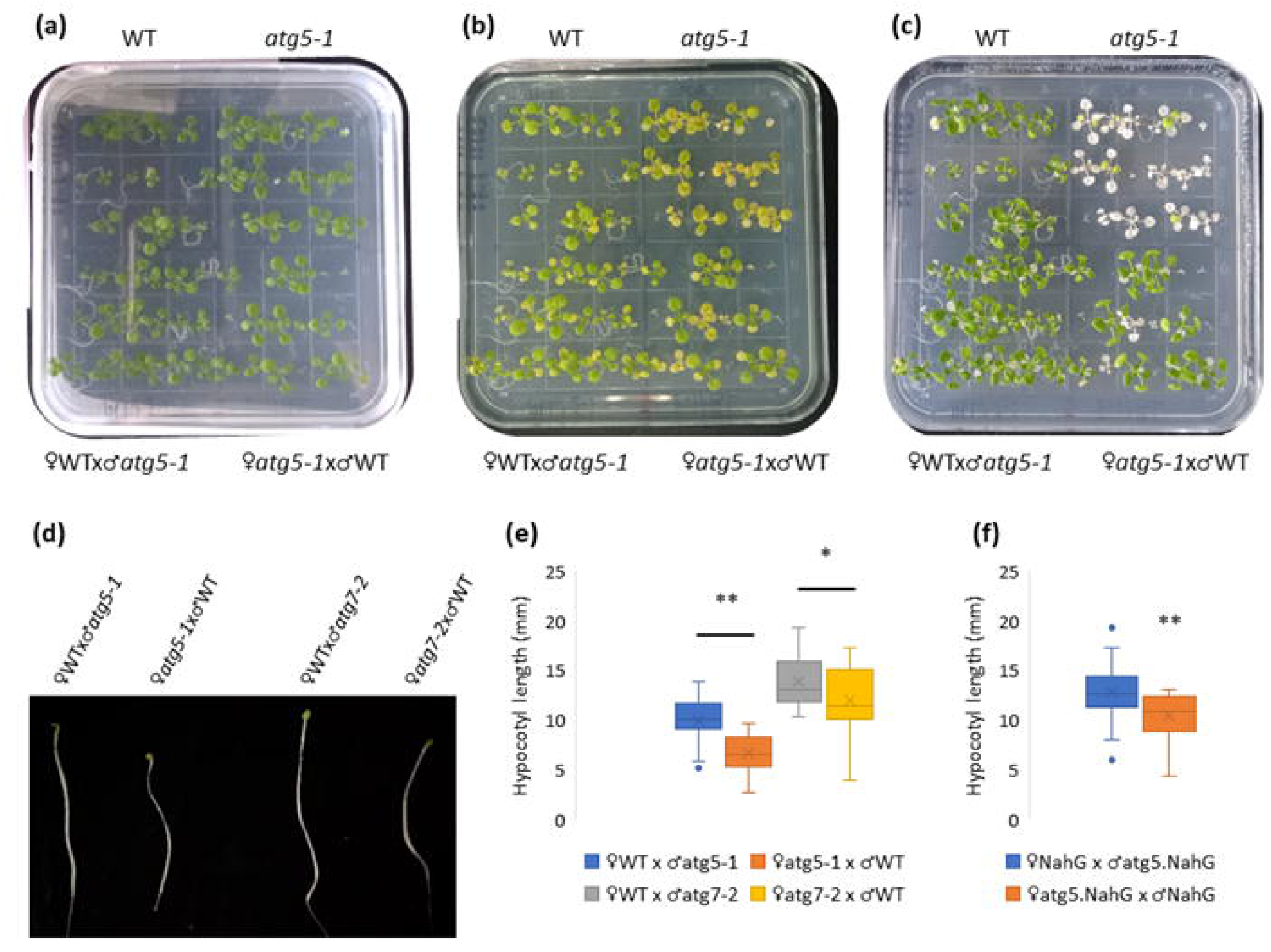
F1 seedlings display a WT under carbon starvation and recovery, yet differential growth as etiolated seedlings. (a-c) Representative images of F1 progeny of reciprocal crosses between WT and *atg5-1* and their respective parental lines. Seeds were germinated and grown on Nitsch plated without exogenous sucrose for 14 days after imbibition (a). The plants were incubated in the dark for additional 7 days (b). The plants were then returned to light for another 7 days to examine their recovery (c). The experiment was performed in triplicate. (d) Representative image of F1 progeny of reciprocal crosses between WT and *atg5-1* or *atg7-2*. Seeds were sown on Nitsch plates without exogenous sugars and grown in the dark for 7 days after imbibition. (e) Hypocotyl length of the lines described in (d), as quantified by ImageJ. Data are presented as a box&whiskers plot. One asterisk (p<0.05) or two asterisks (p<0.01) denote a significant difference following Student’s t-test (n=15-30). (f) Hypocotyl length of F1 progeny of reciprocal crosses between NahG and *atg5*.NahG. Seeds were sown on Nitsch plates without exogenous sugars, grown in the dark for 7 days after imbibition, and their hypocotyl length was measured by ImageJ. Two asterisks (p<0.01) denote a significant difference following Student’s *t-test* (n=20-30).

*atg* mutants display an early senescence phenotype (Marshall and Vierstra, 2018), which was shown to stem from an accumulation of salicylic acid during plant aging (Yoshimoto et al., 2009). We thus wished to ascertain whether the observed phenotype did not originate from the premature senescence of the mother plant, resulting in diminished reserve accumulation. To that end, we performed reciprocal crosses between NahG and *atg5*.NahG plants. NahG is a bacterial enzyme that degrades salicylic acid, and its overexpression in an *atg* mutant background inhibits the early senescence phenotype (Yoshimoto et al., 2009). F1 etiolated seedlings from maternal *atg5*.NahG plants displayed shorter hypocotyls than seedlings from maternal NahG plants, similar to F1 seedlings from maternal *atg* mutant origin (Fig. 3f). This result suggests the phenotype observed for F1 seedlings from maternal *atg* mutants is not a result of early senescence.

### F1 seeds of maternal *atg* mutant plants display reduced storage protein but no difference in lipid content

We next wished to test our hypothesis of reduced storage compounds in F1 seeds of maternal *atg* mutant origin. As Arabidopsis is an oil seed, we performed lipidomics of dry F1 seeds of reciprocal crosses between WT, *atg5-1,* or *atg7-2*. Though we annotated more than 120 lipids, we did not observe any separation between the groups in the principal component analysis (PCA – Fig. 4a). A closer look at TAG species revealed no significant differences or apparent trends between F1 seeds of WT and *atg* mutant maternal origin (Fig. 4b). These results are in line with previous works, demonstrating no reduction in the total lipid content of *atg* mutant seeds compared to WT seeds (Minina et al., 2018 1218).

**Fig. 4:**
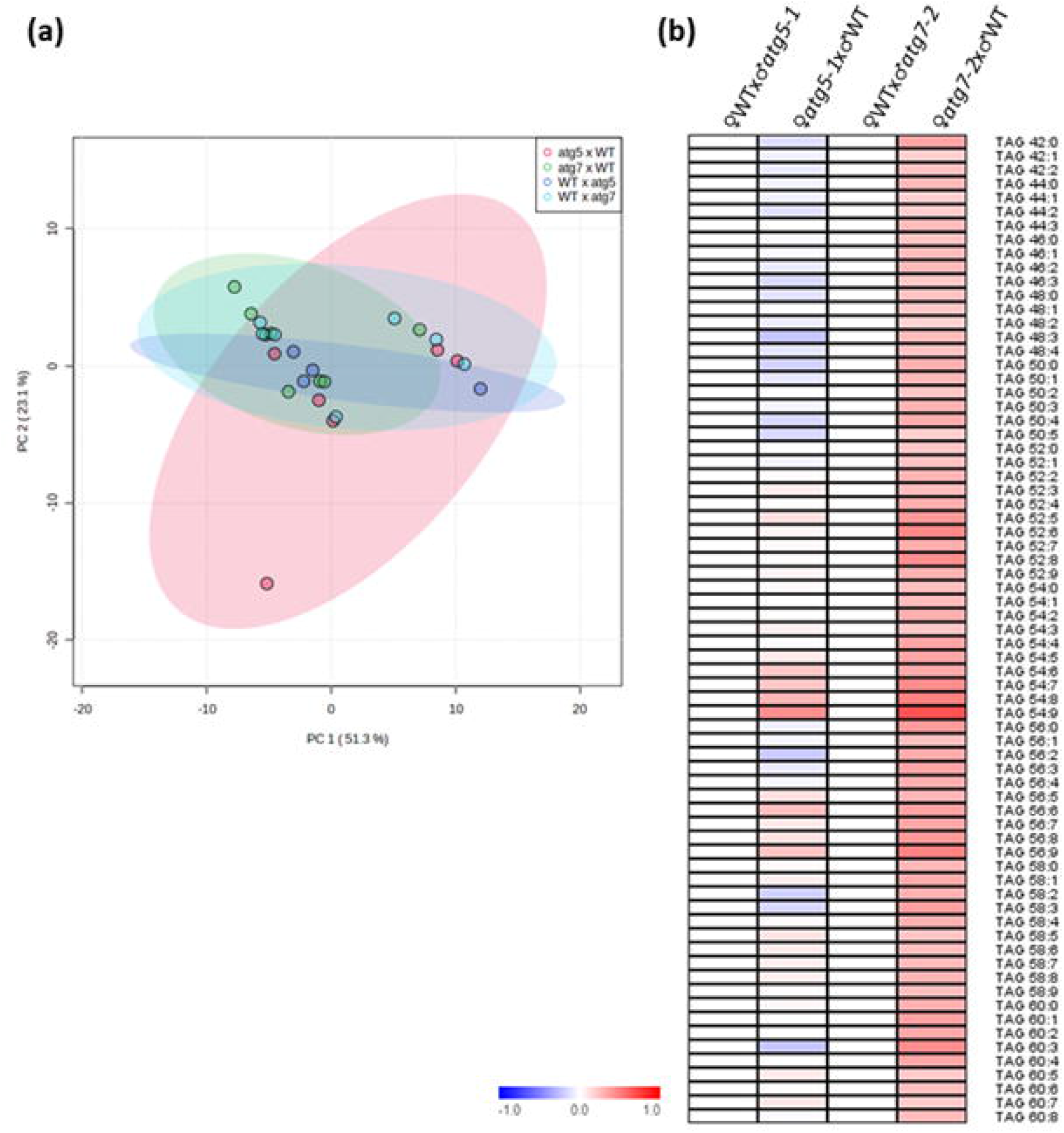
Autophagy deficiency in the mother plant does not affect seed lipid content. Dry seeds of F1 progeny of reciprocal crosses between WT and *atg5-1* or *atg7-2* were collected, and their lipid content was analyzed by UPLC-MS (n=4-8). Detailed results are presented in Supplemental table S1. (a) PCA of lipid levels. (b). Heat map of log2 of TAG relative levels in comparison to seeds from maternal WT lines. No significant differences between reciprocally crossed lines were identified by student’s *t-test*.

We then examined the total protein content of F1 seeds, as they are mostly composed of storage proteins. We extracted total proteins of similar amounts (2mg) of F1 dry seeds. Surprisingly, though no difference in the lipid content was observed, F1 seeds of maternal autophagy-deficient origin contained lower total protein levels than the reciprocally-crossed seeds. The difference was significant for the *atg7-2* crosses, and a similar trend was observed for the *atg5-1* crosses (Fig. 5a). The results indicate that the protein content, but not the lipid content of the seed, is affected by the mother plant, suggesting a possible differential regulation of carbon and nitrogen allocation by autophagy in the mother plant. To confirm our hypothesis, we analyzed the polar metabolites of dry F1 seeds by GC-MS. We could detect 17 metabolites, yet we could not observe significant differences between the reciprocally-crossed lines, including sucrose (Fig. S2). Unfortunately, we did not detect free amino acids and thus could not determine the amounts of glutamine and asparagine in the F1 seeds.

**Fig. 5:**
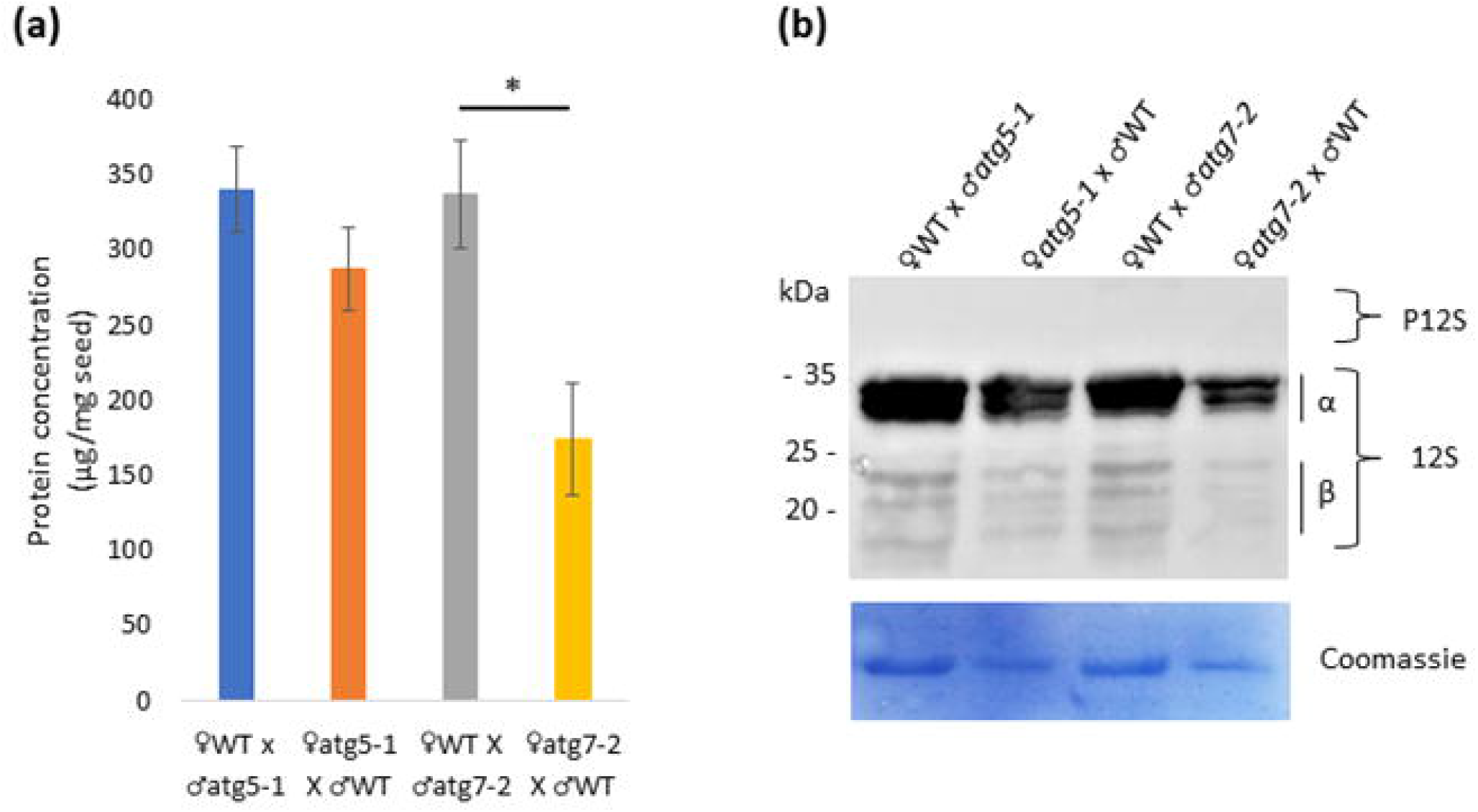
Lack of autophagy in the mother plant alters total seed protein content but not storage-protein processing. Total proteins were extracted from similar amounts of dry seeds of F1 progeny of reciprocal crosses between WT and *atg5-1* or *atg7-2*. (a) Total protein quantification. Data are presented as average ± SE. An asterisk denotes a significant difference between reciprocally crossed lines in Student’s *t-test* (p<0.05, n=3). (b) Total proteins were separated by SDS-PAGE. Top panel – Western blot using anti-12S antibody. Bottom panel – Coomassie blue stain. Three biological replicates were performed. P12S – precursors of the 12S protein, α and β – 12S globulin subunits.

Previous work indicated that homozygous *atg* mutant seeds display altered storage protein processing (Di Berardino et al., 2018). We, therefore, wanted to examine whether the altered protein amounts in F1 seeds correlated with altered storage protein processing. To that end, we performed Western blot analysis against 12S globulin using total proteins extracted from equal seed quantity. Though we observed a general reduction in protein amounts, we did not observe a change in the storage protein pattern (Fig. 5b). For example, no accumulation of globulin precursor (P12S) was observed in F1 seeds, as previously described for *atg5* mutants (Di Berardino et al., 2018). Our results indicate no difference in protein processing in F1 seeds, suggesting that the altered storage-protein processing in *atg* mutant seeds is due to a lack of autophagic activity in the embryo. This is in line with the normal autophagy phenotype of F1 plants (Fi. 3a-c, Fig. S1c-e). We thus show that the reduced protein quality of *atg* mutant seeds stems from a lack of autophagy in the source tissues, while storage protein processing results from autophagy in the sink tissue.

### F1 seeds of maternal *atg* mutant plants display faster germination rates stemming from altered seed coat morphology

A typical phenotype of *atg* mutants is slightly delayed germination and seedling establishment (Avin-Wittenberg et al., 2015). We wanted to determine whether this phenotype stems from autophagy in the mother plant and thus tested germination and seedling establishment rates. We expected no difference in germination or slightly slower germination in the F1 seeds from maternal *atg* mutant origin. Surprisingly, increased germination rates were observed for F1 progeny of *atg* mutant mother plants than WT mother plants (Fig. 6a). No significant differences were seen in seedling establishment of F1 seeds (Fig. 6b). We repeated the experiment for F1 seeds of reciprocally-crossed NahG and *atg5*.NahG plants and received similar results, suggesting the phenotype is unrelated to early senescence (Fig. S3).

**Fig. 6:**
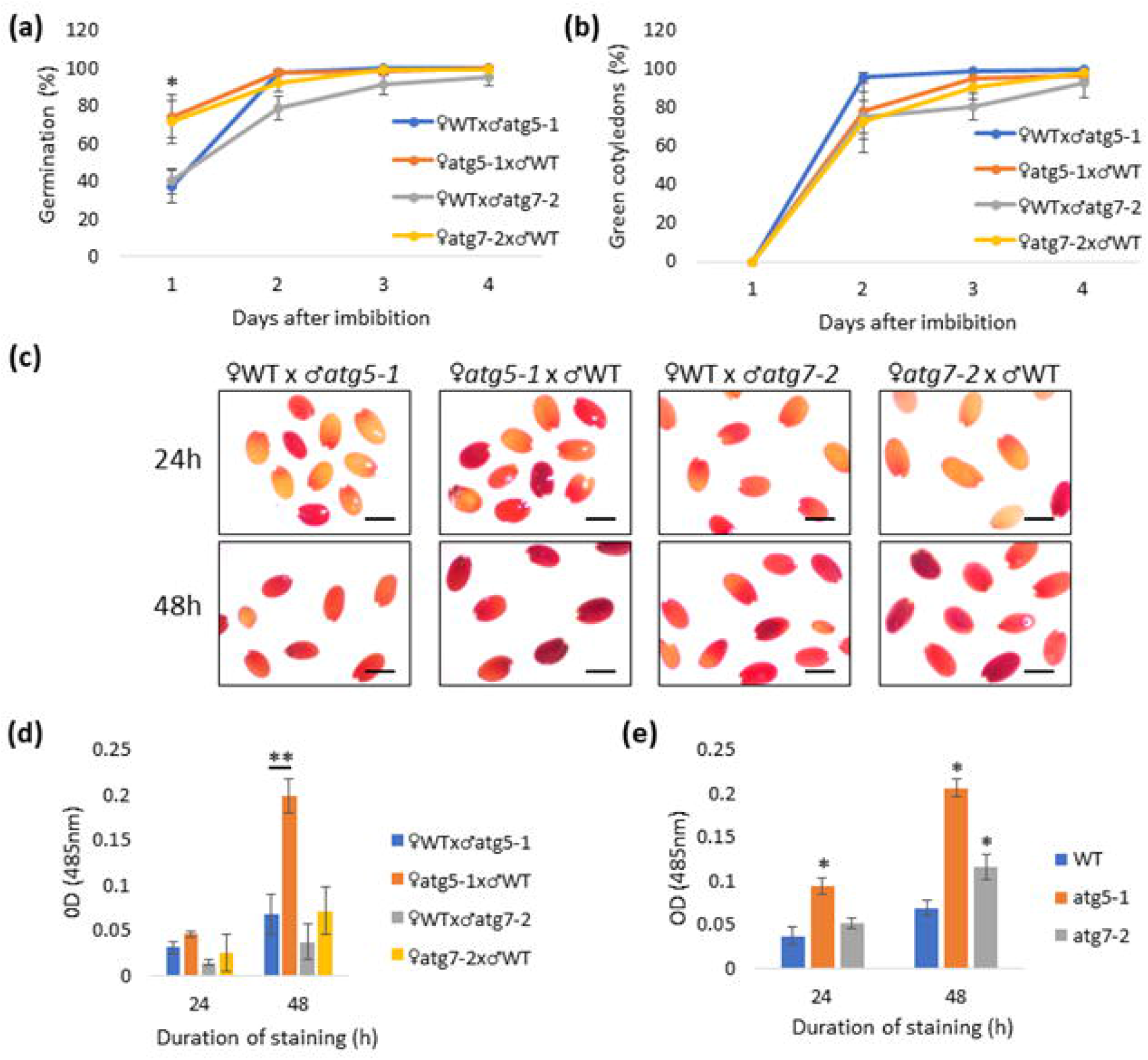
F1 seeds of maternal atg mutants exhibit faster germination and increased permeability. (a,b) F1 seeds of reciprocal crosses between WT and atg mutants were sown on Nitsch plates without exogenous sucrose, imbibed for 72h, and transferred to continuous light conditions. Germination (defined by radicle protrusion) and seedling establishment (defined by the appearance of two green cotyledons) were scored each day for 4 days. (a) Average percent germination is presented ± SE. (b) Average percent of seedling establishment is presented ± SE. An asterisk denotes a significant difference between reciprocally-crossed lines in Student’s *t-test* (p<0.05, n=5). (c) Representative image of F1 seeds from reciprocal crosses after incubation for 24h (top panel) or 48h (bottom panel) in tetrazolium chloride solution. Scale bar=0.5mm. (d, e) Quantification of tetrazolium chloride staining, Stained seeds were ground and dissolved in 95% EtOH. absorption in 485nm was measured. Average absorption is presented ± SE for F1 seeds (d) or homozygous parental lines. One asterisk (p<0.05) or two asterisks (p<0.01) denote a significant difference between reciprocally-crossed lines following Student’s *t-test* (n=5).

We postulated the higher germination rate of F1 progeny of *atg* mutant mother plants resulted from faster water penetration into the seed. We thus examined seed permeability by tetrazolium chloride staining. Tetrazolium salts are amphipathic cations. After penetrating the dead cells of the seed coat, they are reduced by active dehydrogenases (NADH-dependent reductases) in the embryo to red-colored insoluble precipitates composed of formazans (Berridge, 1996). The intensity of red coloration is directly proportional to the permeability of the seeds. We stained dry F1 seeds for 24 and 48 hours. As shown in Fig. 6c and d, F1 seeds of maternal *atg5-1* plants displayed significantly higher water permeability than F1 seeds of maternal WT origin. A similar, albeit insignificant trend, was observed for the *atg7-2* reciprocal crosses. Interestingly, we also observed increased water permeability in homozygous *atg* mutant seeds, suggesting this phenotype is ubiquitous in *atg* mutants (Fig. 6e).

As water permeability is influenced by the seed coat, which is of maternal origin (Marbach and Mayer, 1974), we took a closer look at the seed coat of F1 seeds and their parental lines. We first verified that *ATG5* and *ATG7* were expressed in the Arabidopsis seed coat. A previous study performed detailed gene expression analysis in the seed coast at different developmental stages (Dean et al., 2011); the data is available at the eFP browser (https://bar.utoronto.ca/efp_seedcoat/cgi-bin/efpWeb.cgi)(Baxter et al., 2007). Both *ATG5* and *ATG7* were expressed in the seed coat, starting from 3 days post-anthesis (Fig. S4), strengthening our hypothesis that autophagy is functional in the seed coat during seed development.

We then wished to characterize how seed-coat traits differed between WT and *atg* mutant seeds. We first focused on the hilum, the scar left on the seed coat after detachment from the funiculus. In mature Arabidopsis seeds, the hilum is adjacent to the micropyle (where the radicle will emerge) and faces the chalazal pole (Beisson et al., 2007). The hilum contains high levels of suberin, inhibiting the entrance of water and pathogens. We compared the suberin levels of homozygous WT and *atg* mutant seeds by autofluorescence. We did not observe any differences between the lines (Fig. S5a). We next examined the integrity of the cuticle layer that wraps the embryo (Steinbrecher and Leubner-Metzger, 2018). We stained WT and *atg* mutant seeds with toluidine blue (TB), which stains the layers underneath the cuticle when it is damaged. As a positive control, seeds were pre-incubated with EDTA to perforate the coticule. We did not detect any differences between the lines; none were stained, suggesting their cuticle was intact (Fig. S5b). The seed coat contains phenols in many species, inhibiting water entrance (Werker et al., 1979). We stained WT and *atg* mutant seeds with FeCl_3_ to observe phenols. No genotype displayed dark spots as expected in the presence of phenols (Fig. S5c).

Hydrated Arabidopsis seeds are coated by a gelatinous layer called mucilage, which is mainly composed of cell wall polysaccharides (Voiniciuc et al., 2016). It is deposited in the seed coat epidermal cells during seed development in a structure at the center of the cell termed “columella”. The mucilage is maintained in the epidermal cells following cell death and is released from the seed coat during imbibition (Griffiths and North, 2017). Mucilage is rich in pectin, and its architecture can be visualized by ruthenium red (RR) dye. When F1 seeds were dyed with RR staining, we found that seeds from maternal *atg5-1* plants displayed lower mucilage content than their reciprocal F1 seeds. We observed a similar trend for F1 seeds of WT and *atg7-2*, yet not statistically significant (Fig. 7a,b). Adherent mucilage is partially anchored by cellulosic rays (Sullivan et al., 2011). We performed calcofluor staining of F1 seeds to visualize mucilage rays. Seeds of maternal *atg5-1* plants displayed higher ray density than their reciprocal F1 seeds. A similar trend was observed for the *atg7-2* cross (Fig. 7c,d).

**Fig. 7:**
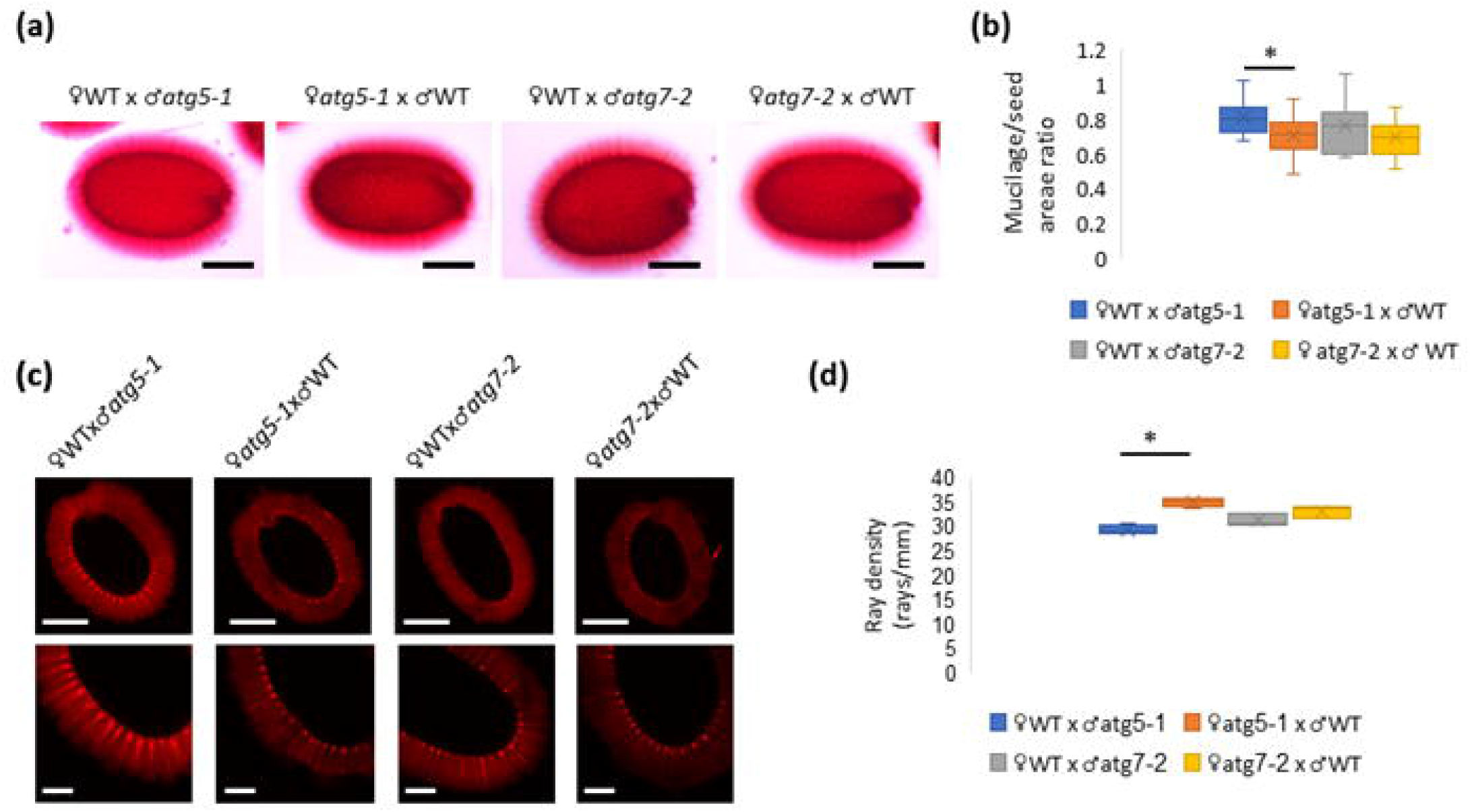
Altered mucilage structure of maternal *atg* mutant F1 seeds. (a,b) F1 seeds of reciprocal crosses between WT and atg mutants were stained with ruthenium red solution. (a) Representative images of F1 seeds from reciprocal crosses. Scale bar = 200µm. (b) Quantification of mucilage area. Data are presented as a box&whiskers plot. One asterisk denotes a significant difference following Student’s *t-test* (p<0.05, n=10). (c,d) Calcofluor staining for β-glucans of F1 seeds. (c) Representative images of whole-seed views (top panel) and close-ups (bottom panel). Scale bars = 250µm (top panel) and 100µm (bottom panel). (d) Quantification of ray density. Data are presented as a box&whiskers plot. One asterisk denotes a significant difference between reciprocally-crossed lines following Student’s *t-test* (p<0.05, n=3-4).

Finally, we assessed the general structure of F1 seeds and seed coat cells by scanning electron microscopy (SEM). F1 progeny of maternal *atg* mutants displayed altered seed shape compared to F1 seeds originating from WT mother plants, stemming from reduced seed width (Fig. 8a-top panel,b). We also observed that the seed coat cells of F1 seeds from *atg* mutant mother plants were misshapen (Fig. 8a-bottom panel). The columella of these cells occupied more of the cell area than in seeds from WT mother plants. When we quantified the ratio between the columella and total cell area, we observed a significantly higher ratio for F1 seeds of maternal *atg7-2* plants than WT plants (Fig. 8c). A similar trend was seen in F1 seeds of *atg5-1* and WT reciprocal crosses (Fig. S6). We thus postulate the increased water permeability of maternal *atg* mutants stems from altered columella development, leading to differential mucilage accumulation.

**Fig. 8:**
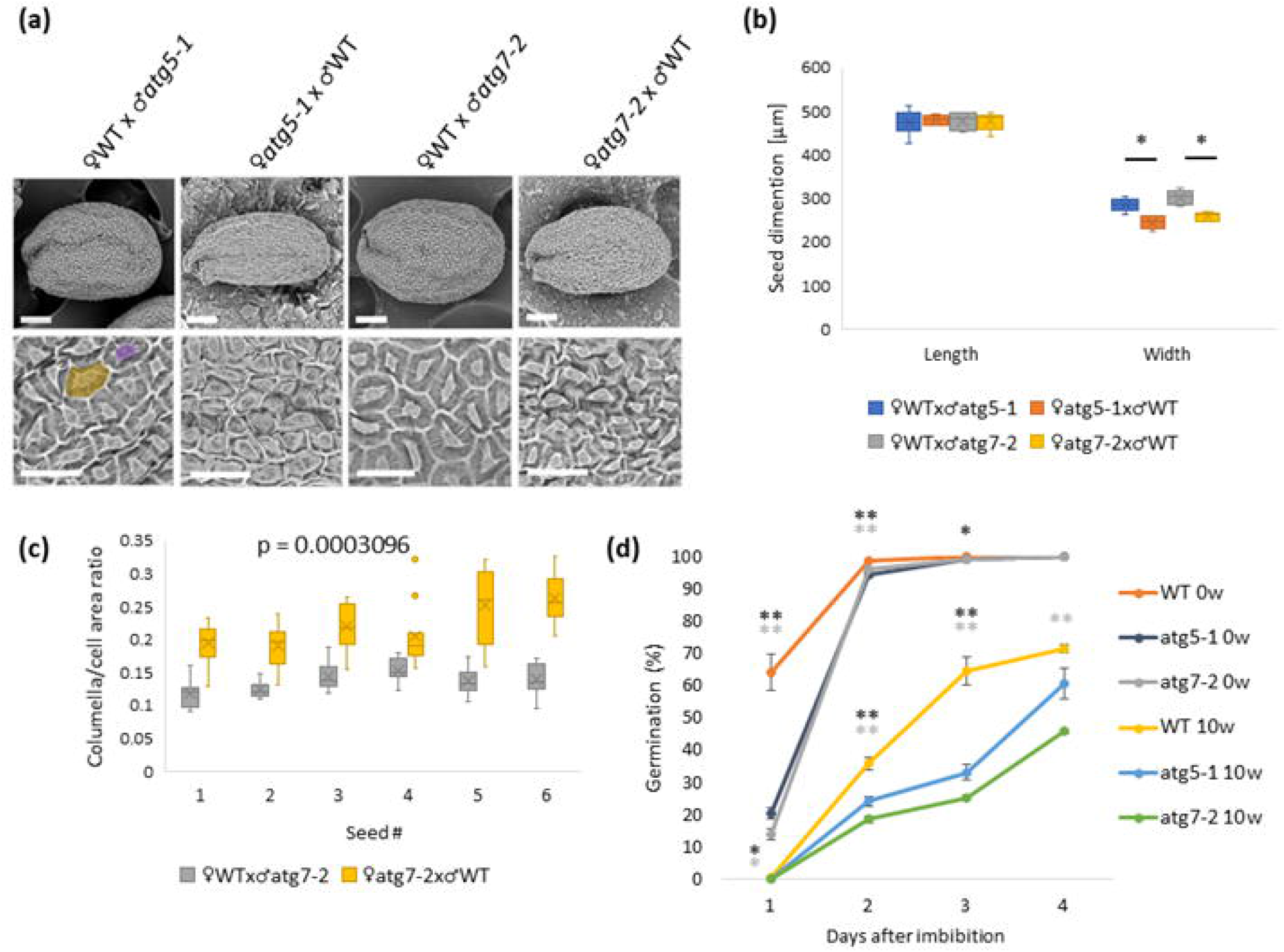
Autophagy in the mother plants has a negative effect on seed shape and aging. (a) Scanning electron microscopy (SEM) of F1 seeds produced by reciprocal crosses between WT and *atg* mutant plants. Top panel, scale bar = 100µm, bottom panel, scale bar = 50µm. The columella tip is marked in purple, and the cell area is marked in yellow (bottom panel). (b) Seed dimensions were measured by ImageJ. Data are presented as a box&whiskers plot. One asterisk denotes a significant difference between reciprocally-crossed lines following Student’s *t-test* (p<0.05, n=6). (c) The ratio between the columella and total cell area in individual F1 seeds of WT and *atg7-2* reciprocal crosses. Data are presented as a box&whiskers plot, p-value denotes the statistical difference between the seeds by mixed model analysis (n=12-15 cells per seed). (d) WT and *atg* mutant seeds without artificial aging and following 10 weeks of artificial aging were sown on Nitsch plates without exogenous sucrose, imbibed for 72h, and transferred to continuous light conditions. Germination (defined by radicle protrusion) was scored each day for 4 days. Average percent germination is presented ± SE. One asterisk (p<0.05) or two asterisks (p<0.01) denote a significant difference from WT at each time point following Student’s *t-test* (n=5, dark gray - *atg5-1*, light gray – *atg7-2*).

### *atg* mutants exhibit more rapid seed aging compared to WT plants

The seed coat structure, as well as seed permeability, play a vital role in seed longevity and aging (Zhou et al., 2019). We thus hypothesized that *atg* mutant seeds would be more sensitive to seed aging. We performed artificial aging of WT and *atg* mutant seeds and tested their germination ability following aging. Without aging, *atg* mutants displayed slightly delayed germination, as previously described (Avin-Wittenberg et al., 2015). Indeed, all lines were affected by seed aging, showing reduced germination after 10 weeks of artificial aging. As expected, the germination of *atg* mutants was more strongly affected by artificial aging than that of WT seeds. This observation strengthens the link between the altered seed coat of *atg* mutants and seed longevity (Fig. 8d). We also scored seedling development following 5 weeks of artificial aging (Boyes et al., 2001) to assess the longer-term effects of seed aging. Artificial aging resulted in delayed development of all the lines. Yet, *atg* mutants developed slower than WT seedlings, further strengthening our assumption that their increased water permeability results in reduced seed longevity (Fig. S7).

## Discussion

Autophagy is gaining attention as a fundamental mechanism in plants, affecting differentiation (Rodriguez et al., 2020), stress response (Tang and Bassham, 2021), and nutrient availability and remobilization (Magen et al., 2022). However, the standard approach to studying plant autophagy involves using ubiquitous knockout or knockdown mutants. Though applicable in many cases, it may obscure our understanding of the function of autophagy in specific tissues or time points, specifically when examining source-sink relationships. *atg* mutant seeds display a plethora of phenotypes, from reduced storage protein processing (Di Berardino et al., 2018) to altered C/N ratio (Guiboileau et al., 2012). Yet, it was impossible to determine the tissue responsible for the phenotype using the current system.

We implemented a novel approach to address this issue described above, performing reciprocal crosses to separate autophagy in the mother plant and the embryo (Fig. 1). This allowed us to determine the role of autophagy exclusively in the mother plant during seed development. We used two well-studied *atg* mutants, *atg5-1* (Yoshimoto et al., 2009) and *atg7-2* (Hofius et al., 2009), to come to general conclusions regarding autophagy function. Although some phenotypes were significant only for one *atg* mutant line, the other line used always showed a similar trend. This aligns with previous studies, in which the phenotypes of different *atg* mutants were not always at the same degree of severity yet displayed a similar trend (Barros et al., 2017; Minina et al., 2018).

The main challenge of our approach is the relatively small number of seeds we could achieve by reciprocal crosses. We were thus unable to perform certain analyses requiring larger material quantities, such as seed coat composition and metabolic flux analysis. Generating transgenic plants in which autophagy is selectively downregulated either in the embryo or the mother plant will allow conducting experiments on a larger scale as well as investigate both sides of the “source-sink coin” concerning autophagy activity. In addition, a transgenic system could be used to investigate the seed set phenotype of *atg* mutants, which reciprocal crosses can not address. A recent example is a study examining autophagy in root cap cells that used tissue-specific CRISPR to down-regulate autophagy specifically in this tissue (Feng et al., 2022). Nevertheless, we believe our work lays the foundation for the in-depth investigation of autophagy in discrete plant tissues and the relationship between tissues.

We determined that F1 seeds displayed a WT carbon starvation phenotype when exposed to dark treatment at the seedling stage, suggesting the autophagy mechanism is active in F1 seedlings (Fig. 2a-c, S1c-e). In contrast, when we examined etiolated seedlings grown without exogenous sugars, F1 seeds from maternal *atg* mutants displayed reduced growth (Fig. 2d,e). These results suggest alerted storage compound accumulation, depending on the presence of autophagy in the mother plant. We observed no significant difference in seed lipid content between F1 seeds (Fig. 3). This aligns with previously reported data, in which homozygous *atg* mutant seeds did not display altered total fatty acid content. Interestingly, the same paper showed that overexpression of *ATG5* or *ATG7* yielded higher fatty acid content (Minina et al., 2018). This could stem from higher sink strength in the mother plant, pushing carbon to the seeds, or increased demand by the embryo.

Unlike lipids, we observed reduced protein levels in F1 seeds of maternal *atg* mutant origin (Fig. 4a). Previous studies showed that *atg* mutant plants remobilize less nitrogen from the mother plant to developing seeds (Guiboileau et al., 2012; Li et al., 2015). Together with our observations, this strengthens our hypothesis that the altered protein amounts in F1 seeds stem from differential remobilization from the mother plant. Surprisingly, carbon remobilization (depicted by the lipid quantity) is less influenced by autophagy than nitrogen remobilization (depicted by the protein quantity). Interestingly, we did not observe any difference in storage protein processing, as described for *atg5* mutant seeds (Di Berardino et al., 2018), suggesting that autophagy in the embryo functions in storage protein processing. The involvement of autophagy in storage protein delivery in seeds has been implied yet hardly investigated, and mostly in monocot, rather than dicot seeds (Levanony et al., 1992; Reyes et al., 2011). More work is needed to elucidate the role of autophagy in storage protein processing, and we believe downregulating autophagy specifically in the embryo will shed light on this fascinating topic.

An important seed tissue we did not address in the scope of this research is the endosperm. This is a triploid tissue that is partly maternal (2n) and partly paternal (1n) (Bentsink and Koornneef, 2008). This uneven distribution creates a situation in which F1 seeds of maternal WT plants possess more copies of *ATG* genes in their endosperm than F1 seeds of maternal *atg* mutant origin (Fig. 1). We could not directly investigate this issue due to the seed amount issue described above. We are unaware of studies examining the effect of *ATG* gene copy number on plant autophagy activation. However, We did observe that F1 seedlings, containing a lower copy number of *ATG* genes, behaved similarly to WT plants under carbon starvation (Fig. 2a-c, S1c-e), suggesting that one copy is sufficient for autophagy activation. Nonetheless, we cannot disregard the contribution of the endosperm to proper embryo growth and development (Song et al., 2021). Future studies are necessary to investigate the function of autophagy in the seed endosperm, perhaps utilizing monocot seeds that have a larger endosperm.

We were most surprised by the observation that F1 seeds of maternal *atg* mutant plants exhibited faster germination compared to F1 seeds from maternal WT plants (Fig. 5a). This was unexpected, as homozygous *atg* mutant seeds display delayed germination compared to WT seeds (Avin-Wittenberg et al., 2015). Of note, the early germination phenotype does not result from the early senescence phenotype of the *atg* mutant mother plant, as F1 seeds of stay-green *atg* mutants displayed a similar phenotype (Fig. S3). We attributed this phenotype to increased water permeability of F1 seeds of maternal *atg* mutant origin (Fig. 5c,d). Interestingly, homozygous *atg* mutant seeds also exhibited increased water permeability, though they are known to germinate slightly slower than WT seeds (Avin-Wittenberg et al., 2015). We postulate that the slower germination of homozygous *atg* mutants is, in fact, a compound phenotype affected by the lack of autophagy in the seed coat and the embryo. Examining the germination of seeds in which autophagy is selectively downregulated in the embryo might help differentiate between the two options.

The reason for the increased water permeability of *atg* mutants is still unclear. Yet, we could observe several differences in the seed coat of F1 seeds of maternal *atg* mutant origin, specifically mucilage accumulation and columella shape (Fig. 6, Fig. 7a-c). More extensive analysis of the seed coat of *atg* mutants is required to fully elucidate the causes of this phenotype. One possibility is the involvement of autophagy in regulating the programmed cell death of the seed coat cells (Sueldo and van der Hoorn, 2017), resulting in deformed seed coat in *atg* mutants and improper deposition of mucilage. Our hypothesis is strengthened by the fact both *ATG5* and *ATG7* are expressed in the Arabidopsis seed coat from early stages of seed development (Fig. S4).

To conclude, our work presents a novel approach to dissecting the function of autophagy in source and sink tissues and examining their effect at the whole plant level. We have shown, for the first time, that the seed phenotype of *atg* mutants is a compound phenotype resulting from the lack of autophagy in both the mother plant and the embryo. Moreover, we implicated autophagy in the mother plant in nitrogen, but not carbon remobilization. Our work highlights the importance of studying the role of autophagy in higher resolution while keeping in mind the whole-plant context. We believe that the generation of plant lines in which autophagy is down-regulated explicitly in space or time will prove to be a valuable tool to achieve this goal.

## Materials and Methods

### Plant material and growth conditions

*Arabidopsis thaliana* ecotype Columbia (Col-0) was used in this study. The lines used were *atg5-1* (SAIL_129B079)(Yoshimoto et al., 2009), *atg7-2* (GK-655B06) (Hofius et al., 2009), NahG over-expression (NahG) (Yoshimoto et al., 2009) and atg5.NahG (Yoshimoto et al., 2009). Seeds were surface sterilized with Cl_2_ for 2h and sown on Nitsch medium (Duchefa) plates pH 5.8, with 1% sucrose. The seeds were imbibed at 4°C for 72h in the dark. Plants were grown in a growth chamber (Fitoclima S600PLH, Aralab) at 22°C and 50% humidity in constant light (125 µmol photons m^-2^ s^-1^) for 2 weeks. They were then transferred to pots containing Perlite: Vermiculite: Garden mix (1:1:2 v/v) and grown at 22°C under long-day conditions (8h dark / 16h light) with 150 µmol photons m^-2^ s^-1^.

### Plant crosses and genotype validation

We conducted reciprocal crosses between Col-0 and *atg5-1* or *atg7-2*. Seeds were collected 20 days after the cross. PCR analyses were performed for genotype validation. DNA from 10 plants grown from crossed seeds was extracted for each cross (Dellaporta et al., 1983). The *atg5-1* crosses were analyzed with *ATG5* forward primer (ATTCACTTCCTCCTGGTGAAG), and either *ATG5* reverse primer for the WT allele (TTGTGCCTGCAGGATAAGCG) or T-DNA left border for the mutant allele (TAGCATCTGAATTTCATAACCAATCTCGATACAC). The *atg7-2* crosses were analyzed with *ATG7* forward primer (GATTCAATCAACTCGCTAAGGCGT), and either *ATG7* reverse primer for the WT allele (TGCTAATTCCATGGATCCAAC) or T-DNA left border for the mutant allele (CCCATTTGGACGTGAATGTAGACAC). Reciprocal crosses between NahG and atg5.NahG were performed as well.

### Embryo development

Siliques (5 from each genotype) were collected 8 and 12 days after flowering (DAF) and opened. The seeds were discolored for 30min in Hoyer’s Solution (7.5g gum arabic, 100g chloral hydrate, 5ml glycerol, and 30ml water). Discolored seeds (40±5) were then placed between a slide and coverslip and observed using a Nomarski objective (20x/0.50 DIC M/N2) on an Eclipse 80i microscope (Nikon) coupled to a digital camera.

### Carbon starvation and recovery

Reciprocally crossed seeds and seeds of the parental lines were sown on Nitsch plates without exogenous sugars (24 plants per line per plate). After imbibition, the plants were placed in a growth chamber for 2 weeks, covered by aluminum foil for 1 week, and exposed to light for an additional week. The plates were photographed before and after carbon starvation and following a week of recovery.

### Hypocotyl length measurements

Reciprocally crossed seeds were sown on Nitsch plates without exogenous sugars. Following imbibition, covered plates were transferred to a growth chamber, and the seeds were grown vertically in dark conditions for 7 days. The plates were photographed, and hypocotyl length was measured by ImageJ (Schneider et al., 2012)(15-40 seedlings per line).

### Metabolite extraction for lipidomics and GC-MS analysis

10 seeds from reciprocally crossed lines were weighed using the MX5 microbalance (Mattler Toledo) as previously described (Bromke et al., 2015). Lipids and polar metabolites were extracted from 4-8 biological replicates using the MTBE method (Salem et al., 2016). The organic phase was dried and used for lipid analysis, and the polar phase was dried and used for polar metabolite analysis.

### Lipid analysis by liquid chromatography-mass spectrometry (LC-MS)

Vacuum-dried organic phases were processed using ultra-performance liquid chromatography (on a C8 reverse-phase column) coupled with Fourier transform mass spectrometry (Q exactive mass spectrometer; Thermo Fisher Scientific) in positive and negative ionization modes. Processing of chromatograms, peak detection, and integration were performed using REFINER MSH 10 (GeneData). Mass spectrometry data processing included removing fragmentation information, isotopic peaks, and chemical noise. Selected features were annotated using an in-house lipid database (Lapidot-Cohen et al., 2020). More information can be found in Supplemental Table S1.

### Polar metabolite analysis by gas-chromatography-mass spectrometry (GC-MS)

Vacuum-dried polar metabolites were measured by the Agilent 7200B GC/Q-TOF. The injection and separation procedures were performed according to Dahtt *et al.* using the DB-35MS column (Dhatt et al., 2019). Metabolite detection and annotation were performed by Quant software (Agilent), according to an accurate mass library of known markers generated by our group. More information can be found in Supplemental Table S2. Following blank subtraction, the peak area of each metabolite was normalized to the internal standard (i.e., ribitol) and the weight of the sample.

### Total protein extraction, quantification, and immunoblotting

Seeds (2mg, three replicates for each line) were ground. Total proteins were extracted using a protein extraction buffer [50mM pH 7.5 Tris-HCl, 3% SDS, 10% glycerol, 150mM NaCl, 1 × protease inhibitor (Roche), 20mM DTT], based on a previously described extraction buffer (Di Berardino et al., 2018). After the addition of buffer and centrifugation for 15min at 13,000g, supernatants were collected. Samples were incubated in SDS-PAGE sample buffer (Laemmli, 1970) for 10min at 100°C.

Total protein content was measured based on Minamide and Bamburg method(Minamide and Bamburg, 1990)with slight modifications. Bovine serum albumin (BSA) powder dissolved in distilled water to a concentration of 10 mg/ml was used to generate a standard curve, and a series of dilutions were made (0, 0.156, 0.312, 0.625, 1.25 and 2.5mg/ml). A grid of 1.5×1.5 cm squares was drawn on a sheet of Whatman number 3 filter paper and divided into 24 squares. 10µl of the standard curve dilutions and the total protein samples were applied within the squares and stained in Coomassie brilliant blue (0.25% w/v in destain solution: 50% methanol, 10% acetic acid, 40% DDW, Raymond A. Lamb) for 10 minutes.

The Whatman paper was destined with destain solution (50% methanol, 10% acetic acid, 40% distilled water) for 10 minutes and washed in water for 2 minutes. Samples were transferred to a 24-well plate with 200µl DMSO in each well. The plate was mixed gently for 10 minutes. 100µl of each sample was taken to a new 96-well plate in two technical replicates, and the absorbance of the standard curve dilutions and samples were obtained at 600 and 400 nm.

For immunoblotting, total proteins were separated by SDS-PAGE using Tris-Glycine running buffer (Fling and Gregerson, 1986) and transferred to a polyvinylidene difluoride membrane (GE Healthcare Life science), as previously described (Towbin et al., 1979). Equal loading was validated using Coomassie Brilliant Blue staining of the membrane. For immunodetection, 12S Globulin antibody (Agrisera; AS204403) was used in combination with horseradish peroxidase-conjugated goat anti-rabbit antibody (Sigma; ab920).

### Seed germination assay

Seeds were sown as described above on Nitsch plates without exogenous sugars (five plates per line, 10-60 seeds from different siliques per plate). After imbibition, the plates were transferred to a growth chamber, and the number of germinated seeds was counted every day for 4 days. Germination was scored by radicle protrusion and seedling establishment by the appearance of two green cotyledons. Germination percentage was calculated accordingly.

### Seed permeability staining

<u>D</u>ry seeds (five replicates of 20 seeds) of each genotype were placed in a 1.5ml microcentrifuge tube. 1ml of 1% (w/v) aqueous solution of 2,3,5-triphenyltetrazolium chloride (Sigma) was added to each tube as previously described (Vishwanath et al., 2014). The tubes were incubated in an air incubator at 30°C in the dark for a period of 24 and 48 hours. After incubation, seeds (10 seeds of each genotype) were observed for changes in seed color and imaged using a stereomicroscope.

The extraction of formazans was performed as previously described (Vishwanath et al., 2014). After incubation, as mentioned above, seeds were washed twice with distilled water. Then 1ml of 95% ethanol was added to the seeds, and the mix was ground using a mortar and pestle. The ground seed material in ethanol solution was transferred to a microcentrifuge tube, and the final volume was adjusted to 1ml with 95% ethanol. The tubes were immediately centrifuged at 15,000xg for 3min. The supernatant was collected into a spectrophotometer cuvette, and the absorbance of the formazan extracts was measured at 485nm using a spectrophotometer. As the blank, 95% ethanol solution was used.

### Seed coat visualization

For seed coat visualization, mature seeds were pre-hydrated with water for 90min and then stained as follows. Ruthenium Red staining for pectins was prepared as previously described (Willats et al., 2001), and hydrated seeds were incubated in 0.01% (w/v) Ruthenium Red (Sigma) for 90min at room temperature. Seeds were washed twice in water and then photographed. Image analysis was performed by ImageJ (Schneider et al., 2012). Calcofluor staining was conducted as previously described (Willats et al., 2001) using 25µg ml^-1^ calcofluor (Fluorescent Brightener 28, Sigma) for 20min at room temperature (5 seeds for each genotype, cesa5 (SALK_118491) as a negative control). Seeds were washed for 2h in water and then photographed. Observations were conducted using the FV-1200 confocal microscope (Olympus), with a 10x/0.4 or 20x/0.75 objective microscope equipped with 405nm (Excitation) and 430-470nm (Emission) laser diode. Image analysis was performed by ImageJ (Schneider et al., 2012).

For seed coat autofluorescence, mature seeds of WT, *atg5-1*, and *atg7-2* were illuminated and observed with an IX81 microscope (Olympus) using DAPI interference and absorption filter (excitation filter, 350/50nm; emission filter, 460/50nm) coupled to a digital camera (Li et al., 2007).

For phenol dyeing, green seeds (12-16 DAF) of WT, *atg5-1*, and *atg7-2* were incubated in FeCl_3_ solution (2% in 95% EtOH (Mace, 1963)) for 5min. 95% EtOH was used as A negative control. For cuticle permeability, green seeds (12-16 DAF) of Col-0, *atg5-1*, and *atg7-2* were hydrated for 10min. Hydrated seeds were incubated in an aqueous solution of 0.05% (w/v) Toluidine-Blue (Sigma) for 2 min and then washed with DDW twice (Mace, 1963). Hydration in 0.5% EDTA solution with three DDW washes was used as a positive control.

### Scanning electron microscopy (SEM)

Dry seeds were mounted on aluminum stubs (Electron Microscopy Sciences), sputter-coated with 10nm of Au/Pd (5% Pd), and viewed using “Quanta 200”, field emission scanning electron microscope (Field Emission Instruments. (6 seeds were examined for each genotype. Measurement of 15 epidermis cells from each seed was performed by ImageJ (Schneider et al., 2012).

### Artificial seed aging

Artificial aging was performed as previously described (Nguyen et al., 2012). 50mg of seeds were stored above a saturated NaCl solution in a closed desiccator (relative humidity of 75%) for 70 days. Germination assay was performed on aged and non-aged seeds as described above. For developmental stage scoring, seeds were artificially aged for 35 days and sown on Nitsch plates without exogenous sugars. The plants were grown as described above for 14 days after imbibition. Seedling development was scored daily as previously described (Boyes et al., 2001).

### Statistical analysis

The experiments were conducted in a random blocks design with three to six biological replicates of each genotype. Data were statistically examined using analysis of variance and tested for significant (* = p<0.05, ** = p<0.01) differences using Student’s *t-test* from an algorithm embedded in Microsoft Excel. Heatmaps were generated using the MultiExperiment Viewer (MeV) freely available software application (Howe et al., 2010). Principle Component Analysis was performed using Metaboanalyst 5.0 (Chong et al., 2019). Mixed model analysis was performed using Prism (GraphPad Software).

## Acknowledgments

We thank Dr. Smadar Harpaz-Saad of the Hebrew University of Jerusalem for her assistance with Arabidopsis seed coat analysis. We also thank Dr. Naomi Melamed-Book from the microscopy core facility at The Hebrew University’s Life Science core facility for her technical support with confocal analysis and Dr. Evgenia Blayvas from the Hebrew University’s Unit for Nanoscience, for technical support with SEM.

## Author Contributions

O.E. designed the research, performed research, analyzed data, and wrote the manuscript. S.W., M.A-A, M.A., Y.S., W.J, L.R, and Y.B performed research and analyzed data. T.A-W designed the research and wrote the manuscript. The project was funded by the Israeli Science Foundation, grant number 1942/19.

## Data availability

All data presented in the manuscript is available in the figures or supplementary material.

## Supporting information

**Fig. S1: F1 seedlings of reciprocal crosses are heterozygous and display a WT phenotype.** a,b: PCR analysis for DNA extracted from F1 seedlings (n=5 from different siliques) and parental lines, with primers for the WT allele and T-DNA insertion. (a) atg5-1, (b) atg7-2. c-e: Representative images of F1 progeny of reciprocal crosses between WT and atg7-2 and their respective parental lines. Seeds were germinated and grown on Nitsch plated without exogenous sucrose for 14 days after imbibition (c). The plants were incubated in the dark for additional 7 days (d). The plants were then returned to light for another 7 days to examine their recovery (e). The experiment was performed in triplicate.

**Fig. S2: No difference between F1 seeds in primary metabolites.** Dry seeds of F1 progeny of reciprocal crosses between WT and *atg5-1* or *atg7-2* were collected, and their polar metabolite content was analyzed by GC-MS (n=4-8). Detailed results are presented in Supplemental Data Set S2. (a) PCA of metabolite levels. (b). Heat map of log2 of metabolite relative levels in comparison to seeds from maternal WT lines. An asterisk denotes a significant difference between reciprocally crossed lines by Student’s *t-test* (p<0.05).

**Fig. S3: The early germination of maternal *atg* mutant seedlings is not salicylic acid-dependent.** (a,b) F1 seeds of reciprocal crosses between NahG and *atg5.*NahG mutants were sown on Nitsch plates without exogenous sucrose, imbibed for 72h, and transferred to continuous light conditions. Germination (defined by radicle protrusion) and seedling establishment (defined by the appearance of two green cotyledons) were scored each day for 4 days. (a) Average percent germination is presented ± SE. (b) Average percent of seedling establishment is presented ± SE. An asterisk denotes a significant difference in Student’s *t-test* (p<0.05, n=5).

**Fig. S4: *ATG5* and *ATG7* are expressed in the seed coat during seed development.** *ATG5* (At5g17290 – left panel) and *ATG7* (At5g45900 – right panel) gene expression in Arabidopsis Col-2 seed coat as measured by Dean *et al*. (2011). Seed coat was sampled at 3,7 and 11 days post-anthesis (DPA), and RNA was analyzed by microarray analysis. Data were extracted from the Arabidopsis seed coat eFP browser.

**Fig. S5: No difference between WT and *atg* mutant seeds in several biochemical factors affecting water permeability.** (a) Seed coat autofluorescence was visualized using DAPI filter. All genotypes show saturation in the hilum region (designated by yellow arrowheads) because of high Suberin content. 30-50 seeds per genotype were examined. Scale bar=200µm. (b) TB staining for cuticle permeability. No genotype shows blue staining as the positive control (pre-incubation in 0.5% EDTA solution). 5 siliques were examined per genotype. Scale bar=0.5mm. (c) FeCl3 staining for phenols. No genotype shows dark spots predicted to appear in the presence of phenols. Negative control with DDW. 5 siliques were examined per genotype. Scale bar=0.5mm.

**Fig. S6: Autophagy in the mother plant has a negative effect columella/cell area ratio.** The ratio between the columella and total cell area in individual F1 seeds of WT and *atg5-1* reciprocal crosses. Data are presented as a box&whiskers plot, p-value denotes the statistical difference between the seeds by mixed model analysis (n=12-15 cells per seed).

**Fig. S7: Artificial aging affects seedling development of *atg* mutants.** WT and *atg* mutant seeds without artificial aging and following 5 weeks of artificial aging were sown on Nitsch plates without exogenous sucrose, imbibed for 72h, and transferred to continuous light conditions. Growth stage progression was scored for seedlings grown vertically. Data represent average ± SE (n35). Asterisk indicates significant differences from WT in each aging time (Student’s t test; p<0.05).

**Supplemental Table S1:** Relative lipid content of reciprocally crossed Arabidopsis thaliana F1 seeds.

**Supplemental Table S2:** Relative polar metabolite content of reciprocally crossed Arabidopsis thaliana F1 seeds.

