## Supplemental figures for "Autophagy in maternal tissues contributes to *Arabidopsis thaliana* seed development"

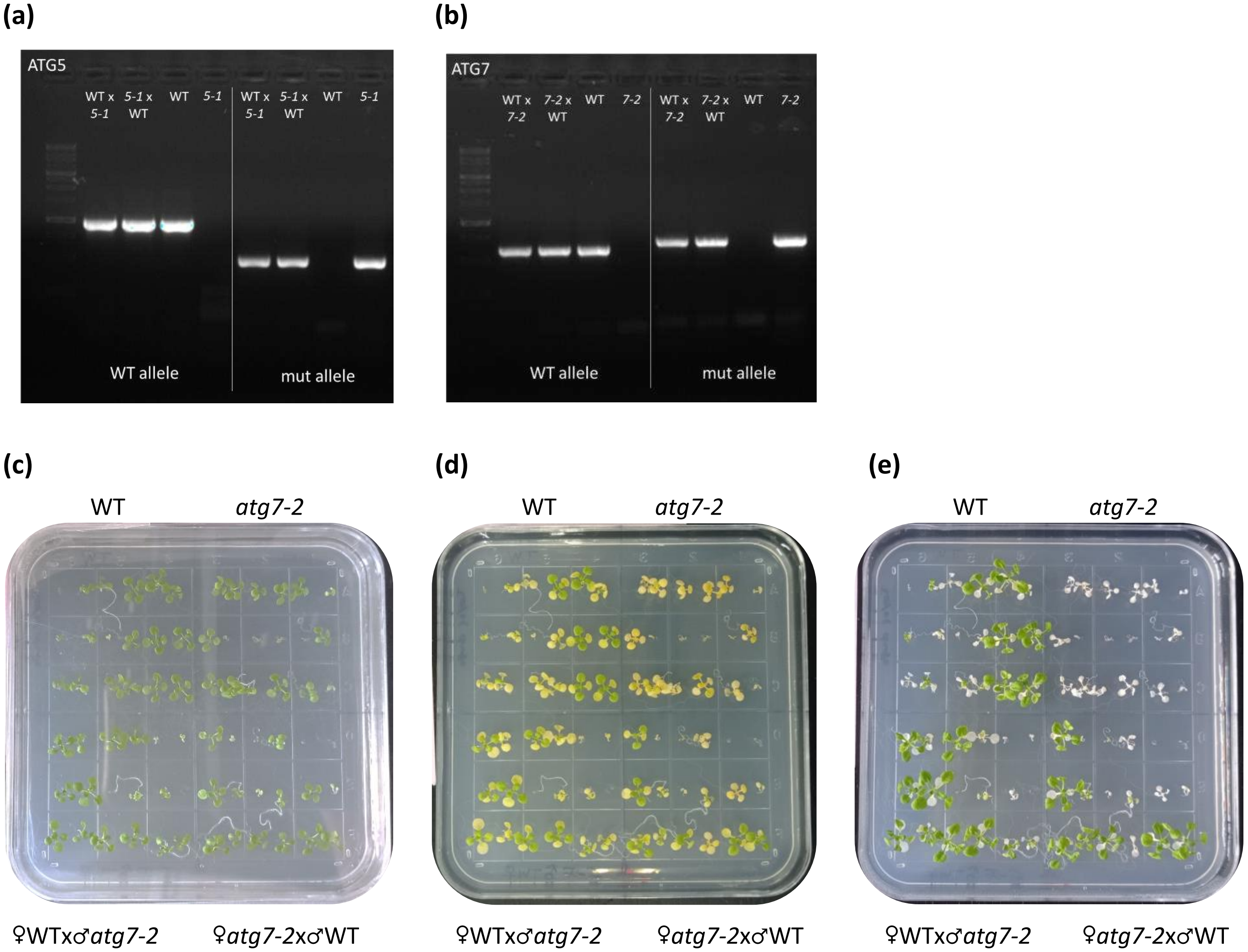


**Fig. S1: F1 seedlings of reciprocal crosses are heterozygous and display a WT phenotype.** a,b: PCR analysis for DNA extracted from F1 seedlings (n=5 from different siliques) and parental lines, with primers for the WT allele and T-DNA insertion. (a) *atg5-1*, (b) *atg7-2*. c-e: Representative images of F1 progeny of reciprocal crosses between WT and *atg7-2* and their respective parental lines. Seeds were germinated and grown on Nitsch plated without exogenous sucrose for 14 days after imbibition (c). The plants were incubated in the dark for additional 7 days (d). The plants were then returned to light for another 7 days to examine their recovery (e). The experiment was performed in triplicate.

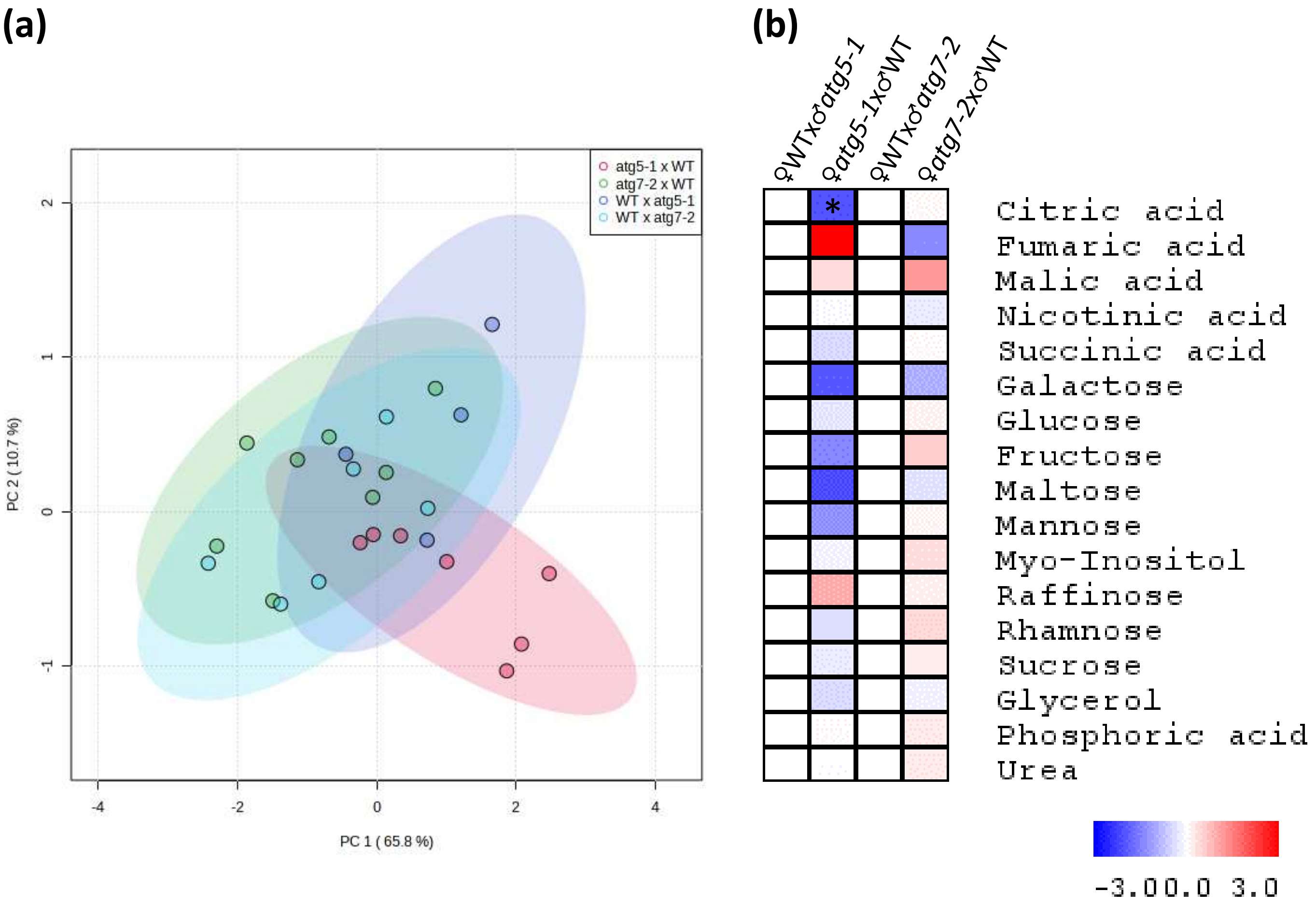


**Fig. S2: No difference between F1 seeds in primary metabolites.** Dry seeds of F1 progeny of reciprocal crosses between WT and *atg5-1* or *atg7-2* were collected, and their polar metabolite content was analyzed by GC-MS (n=4-8). Detailed results are presented in Supplemental Data Set 2. (a) PCA of metabolite levels. (b). Heat map of log2 of metabolite relative levels in comparison to seeds from maternal WT lines. An asterisk denotes a significant difference between reciprocally crossed lines by student's *t-test* ($p<0.05$).

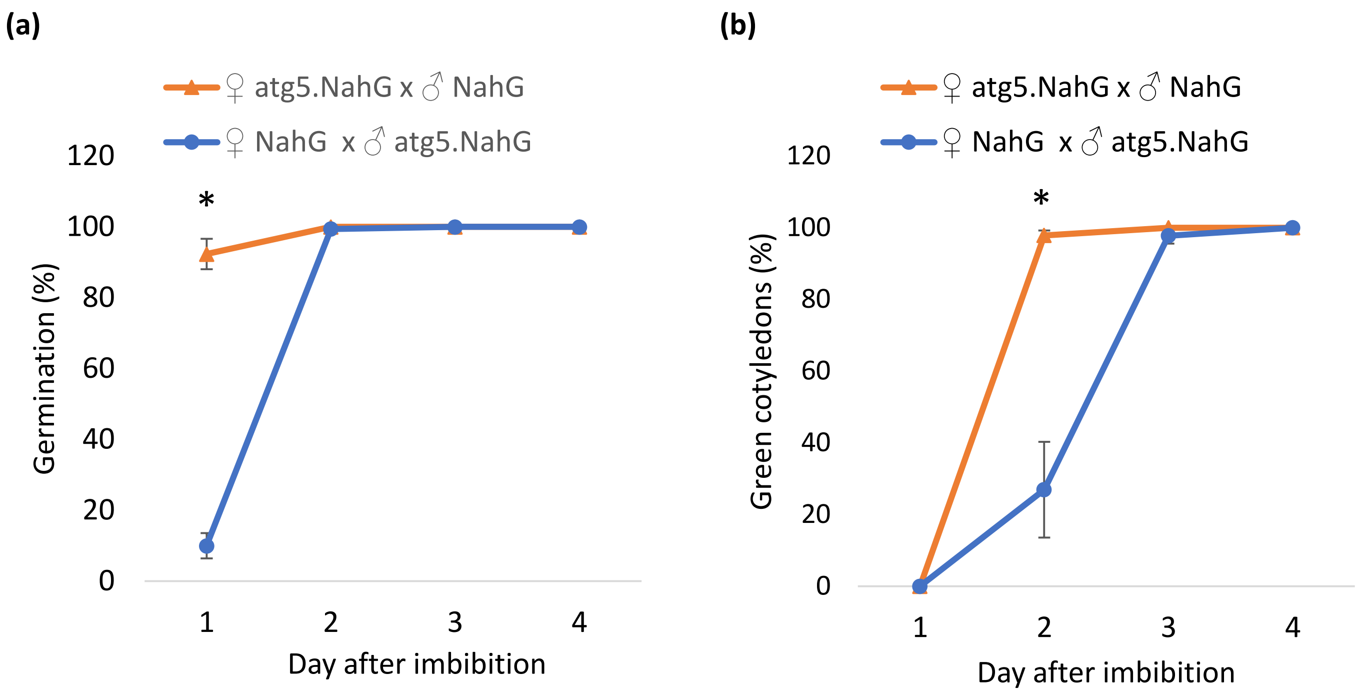


**Fig. S3: The early germination of maternal *atg* mutant seedlings is not salicylic acid-dependent.** (a,b) F1 seeds of reciprocal crosses between NahG and *atg5*.NahG mutants were sown on Nitsch plates without exogenous sucrose, imbibed for 72h, and transferred to continuous light conditions. Germination (defined by radicle protrusion) and seedling establishment (defined by the appearance of two green cotyledons) were scored each day for 4 days. (a) Average percent germination is presented ± SE. (b) Average percent of seedling establishment is presented ± SE. An asterisk denotes a significant difference in student's *t-test* ($p<0.05$, n=5).

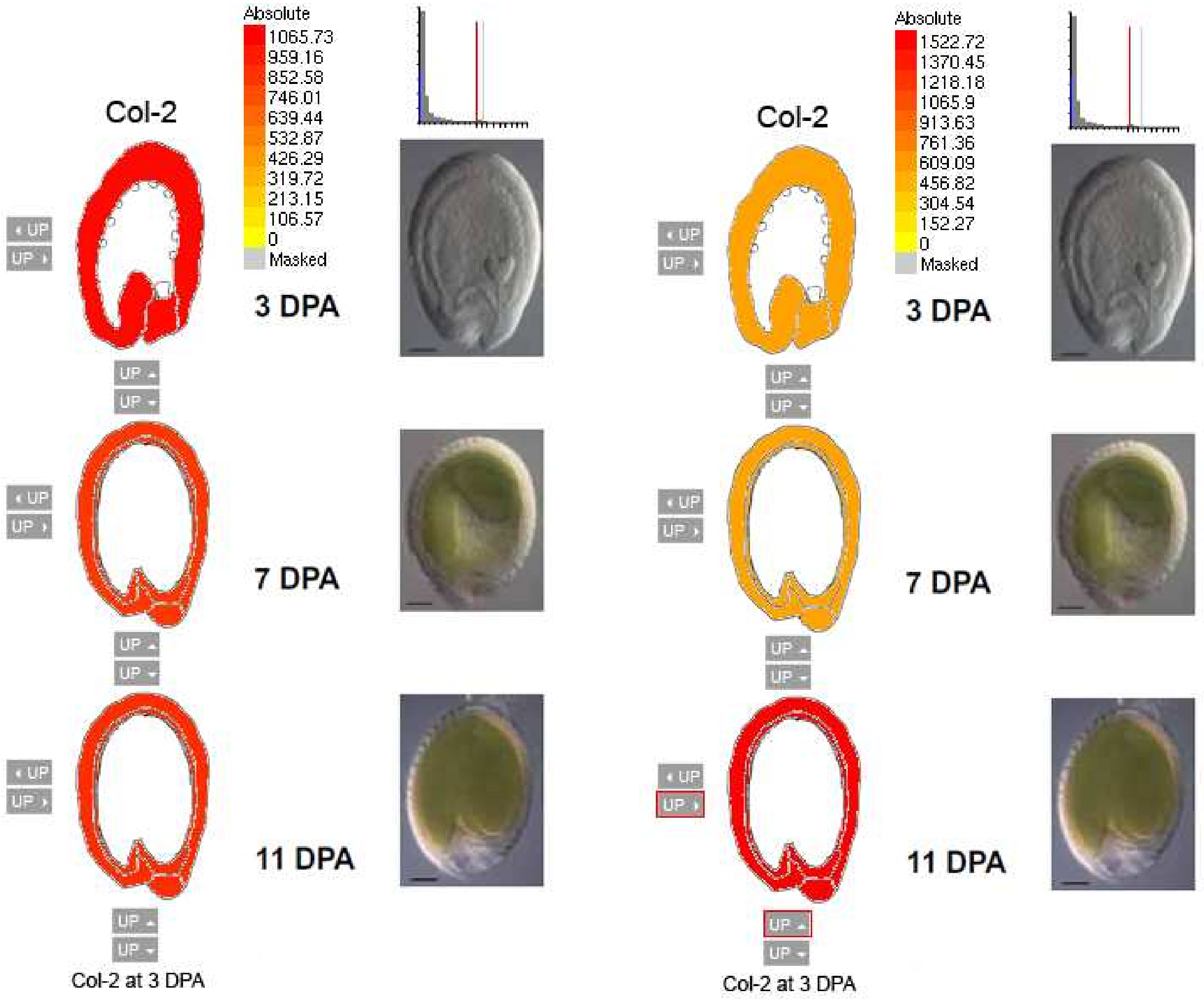


**Fig. S4: *ATG5* and *ATG7* are expressed in the seed coat during seed development.** *ATG5* (At5g17290 – left panel) and *ATG7* (At5g45900 – right panel) gene expression in Arabidopsis Col-2 seed coat as measured by Dean *et al*. (2011). Seed coat was sampled at 3,7 and 11 days post-anthesis (DPA), and RNA was analyzed by microarray analysis. Data were extracted from the Arabidopsis seed coat eFP browser.

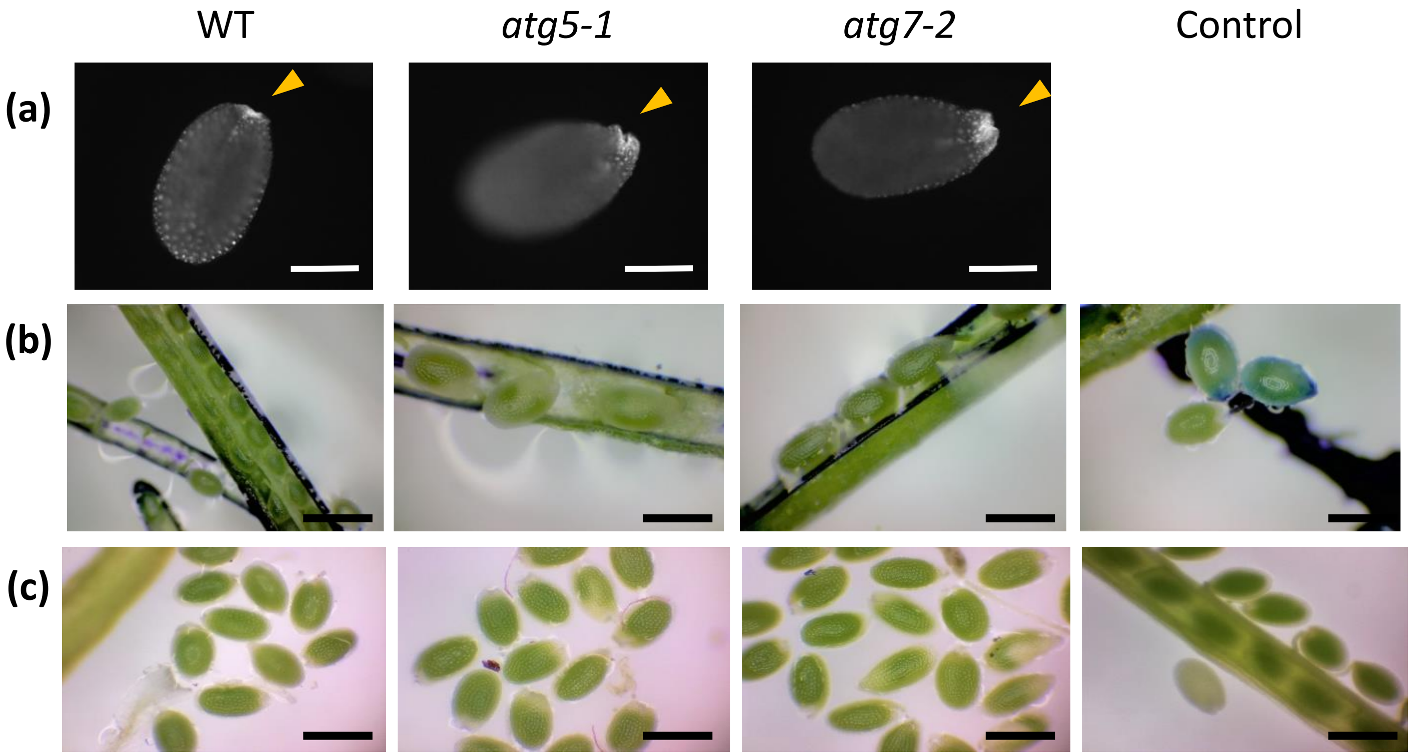


**Fig. S5: No difference between WT and *atg* mutant seeds in several biochemical factors affecting water permeability.** (a) Seed coat autofluorescence was visualized using DAPI filter. All genotypes show saturation in the hilum region (designated by yellow arrowheads) because of high Suberin content. 30-50 seeds per genotype were examined. Scale bar=200µm. (b) TB staining for cuticle permeability. No genotype shows blue staining as the positive control (pre-incubation in 0.5% EDTA solution). 5 siliques were examined per genotype. Scale bar=0.5mm. (c) FeCl3 staining for phenols. No genotype shows dark spots predicted to appear in the presence of phenols. Negative control with DDW. 5 siliques were examined per genotype. Scale bar=0.5mm.

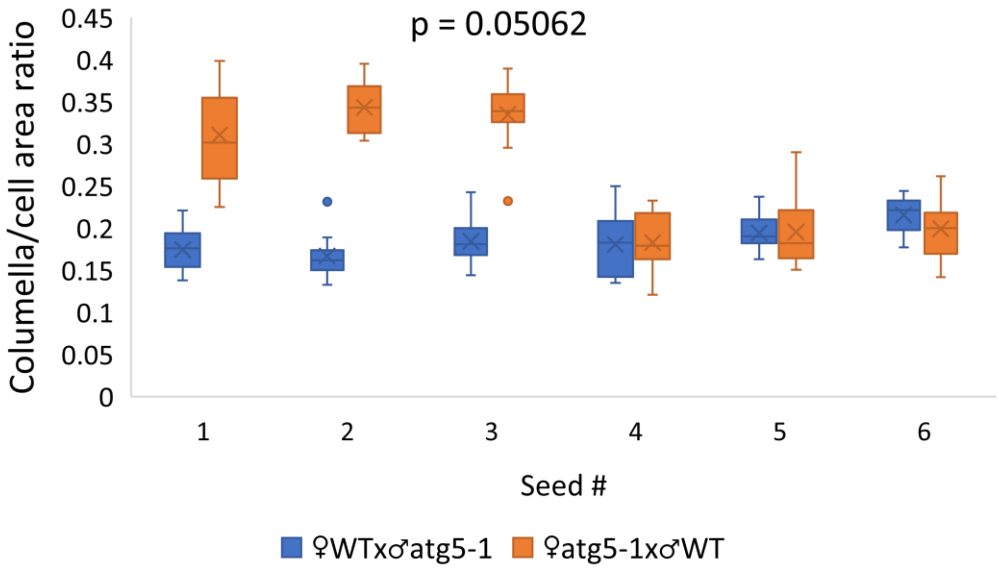


**Fig. S6: Autophagy in the mother plant has a negative effect columella/cell area ratio.** The ratio between the columella and total cell area in individual F1 seeds of WT and *atg5-1* reciprocal crosses. Data are presented as a box&whiskers plot, p-value denotes the statistical difference between the seeds by mixed model analysis (n=12-15 cells per seed).

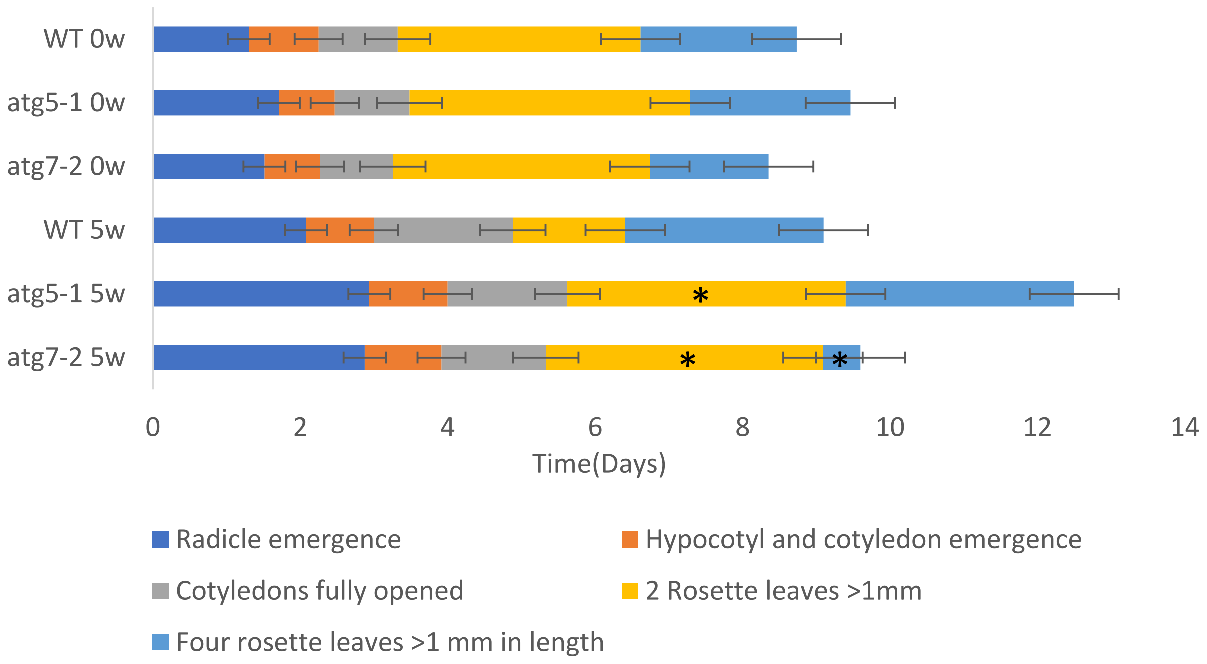


**Fig. S7: Artificial aging affects seedling development of *atg* mutants.** WT and *atg* mutant seeds without artificial aging and following 5 weeks of artificial aging were sown on Nitsch plates without exogenous sucrose, imbibed for 72h, and transferred to continuous light conditions. Growth stage progression was scored for seedlings grown vertically. Data represent average ± SE (n35). Asterisk indicates significant differences from WT in each aging time (Student's t test; $p<0.05$).
